# Polyhyrdoxybutyrate production by *Bacillus marcorestinctum* using a Cheaper substrate and its electrospinned blends with polymer

**DOI:** 10.1101/680124

**Authors:** Swetha Narayankumar, Neethu K. Shaji, Veena gayathri Krishnaswamy

## Abstract

Poly(hydroxybutyric acid) (PHB) and other biodegradable polyesters are promising candidates for the development of environment-friendly and completely biodegradable plastics. One of the major drawbacks in the production of PHB is production costs, since it requires large amount of carbon source. This calls for cheaper substrates that can be used as an alternative carbon source such as agro-industrial residues. In this study, cane molasses was used as an additional carbon source at 2% concentration along with glucose for large scale production of PHB. Ammonium nitrate was used as the nitrogen source and the C:N ratio was maintained at 1:15. The maximum production of PHB was obtained at 24hours of growth which was found to be 0.5g/L and had a dry cell weight of 3.7g/ L.The PHB produced was further analysed by GC-MS Analysis and Transmission Electron Microscopy (TEM).The obtained PHB from scale-up studies were further electrospinned using different blends of polymers.

## INTRODUCTION

Polyhydroxybutyrate (PHB) is a homopolyester belonging to a class of biodegradable intracellular polyesters synthesized by various eubacterial genera and several archea, that are collectively called as Polyhydroxyalkanoates (PHAs). PHB is the best characterized and most extensively studied type amongst the PHAs.^1,2^ They are plastic-like materials whose physical and mechanical features resemble closely to that of conventional petroleum derived plastics such as polyethylene and polystyrene. ^3^ PHB is synthesized by microorganisms as carbon and energy storage compounds within individual cells and its production is induced in a medium containing excess of carbon source along with depletion of an essential nutrient such as nitrogen, phosphorous and oxygen. ^4^

The production of PHB has been most commonly studied on microorganisms belonging to the genera *Alcaligenes, Azotobacter, Bacillus* and *Pseudomonas.* ^5^ The genus *Bacillus* is one of the first Gram-positive bacteria that were identified as being capable of producing PHB.^6^ Since then, it has been widely used in industry and academia due to the stability of its replication and maintenance of plasmids.^7^ However, the higher cost of industrial production of PHB limits the commercial application of bioplastic which means that they have not fully replaced the traditional non-degradable plastics. ^2^ One of the major reasons includes the carbon source which accounts up to 50% of the total production costs. Thus, the use of waste agricultural residues, starch and dairy waste instead of rich medium with glucose can substantially reduce the substrate cost and in turn may even provide value to the waste, thereby downsizing the overall production costs. ^8^

In recent times, Electrospinning has earned much consideration due to its potential for applications in various fields. Electrospun fibers have shown great applicability for novel materials in tissue engineering, wound healing, bioactive molecules delivery as well as sensor, filtration, and nanoelectronic applications. ^9^ This is due to the desirable properties of electrospun fibres which includes increased fibre length with reduced diameter and pore size, large surface area, and high porosity.^10^ This paper demonstrates the effect of cane molasses as an additional carbon source used along with glucose in the production of PHB at the fermentor level. The PHB produced was further characterized by GC-MS and TEM analysis. Also, the PHB produced was electrospinned with different polymer blends

## MATERIALS AND METHODS

### Bacterial Strains

Halotolerant bacterial strains were isolated from Pulicat lake soil, a brackish water lake in TamilNadu. The soil sample was serially diluted and the individual colonies were further subcultured to obtain pure bacterial isolates which were then screened for the production of PHB by primary and secondary screening methods.

Four bacterial strains were found to produce PHB which were biochemically characterized and molecularly identified by 16s rRNA sequencing as *Roseivivax lentus, Bacillus toyonensis, Klebsiella quasipneumoniae subsps.* and *Bacillus marcorestinctum*. The PHB produced by each of the strains was evaluated and the most efficient strain with maximum polymer production was found to be *Bacillus marcorestinctum* which was later labelled as SNKVG 22.The strain was maintained on agar slants and petri dishes and stored at 4°C until use.

### Composition of the culture medium

For PHB scale-up, minimal Salt Medium (MSM) containing the following composition (g/L) – Na_2_HPO_4_ – 4.5, KH_2_PO_4_ – 1.5, MgSO_4_ – 0.2, NaCl – 1, (NH_4_)_2_SO_4_ – 2, CaCl_2_. 2H_2_O – 0.02, NH_4_Fe (III) citrate – 0.05, Glucose – 10 and 1 ml of Trace elements which includes (mg/L) – ZnSO_4_.7H_2_O – 100, H_3_BO_3_ – 300, CoCl2.6H2O – 200, CuSO_4_ – 6, NiCl_2_.6H_2_O – 20, Na_2_MoO_4_.2H_2_O – 30, MnCl_2_.2H_2_O – 25was used.^11^ Carbon source, nitrogen source, Ferric ammonium citrate and Trace elements were sterilized separately at 121°C for 20 min and added to the sterilized medium at the desired concentration. The pH of the medium was maintained at 7.0. Blackstrap molasses were used as an additional carbon source which was also autoclaved separately. The molasses were obtained in 500 ml volume from Dhanyam, Chennai.

### Shake flask study and optimization of growth parameters

Prior to scale-up, a shake flask study was carried out using the efficient bacterial isolate by growing it in 100ml of minimal salt media to evaluate its growth pattern and PHB production for upto 3 days. The optimization for maximum PHB production by the selectedisolate was carried using MSM as the culture media to evaluate several cultural parameters and to determine their effect on growth and PHB production. The optimized value for each parameter was selected and kept constant for further experiments. Several cultural parameterstemperature of incubation (30°C, 35°C and 40°C), and effect of pH (6,7 and 8), carbon:nitrogen ratio (10:1, 15:1, 20:1 and 25:1) and salt concentrations (3%,5% and 7%) were evaluated. The protein content of the cells was evaluated to analyse its growth and PHB content was also determined.

### Scale-up production of PHB

For large scale production, a loopful of culture was taken from a freshly streaked plate and inoculated into a test tube containing 3 ml of nutrient broth. The inoculated tube was incubated at 28°C in an orbital shaker at 200 rpm until an optical density of 2.0 OD_600nm_ was attained. The pre-inoculum culture was transferred into 250 ml of MSM taken in a 500 ml conical flask. It was incubated at 28°C in an orbital shaker at 200 rpm for about 24 hours until a growth of 6.0 – 8.0 OD_600nm_ was attained. The fermentation process was studied in a 3.7 L fermenter(KLF model obtained from Bioengineering AG) with a working volume of 2.5 L using the optimized parameters mentioned above and production medium. Initially, 2500ml of MSM was prepared was and added into the fermenter and the medium was sterilized in situ. Blackstrap molasses was added as an additional carbon source at 2% (v/v) concentration to maintain the overall ratio of carbon and nitrogen sources at 1:15. The entire content of the 250ml flask was then aseptically added to the fermentation medium as the inoculum at 10% concentration. The aeration and agitation were maintained at 1vvm and 500-700 rm respectively. The Dissolved oxygen level was not controlled but was allowed to fall freely until equilibrium level was established. The impeller speed wasmaintained at 500-700 rpm and aeration was set at 1vvm. The pH was maintained at 7.0 using 1 N NaOH and 1 N HCl and throughout the process temperature was kept at 28°C. 0.5 mL of the antifoaming agent, PPG was initially added to the fermenter. About 20 ml culture broth was taken every 4 hours throughout the whole fermentation run and analyzed for cell dry weight (CDW) and PHB yield.

### Cell dry weight estimation

Biomass content of the cells were evaluated by gravimetry. The samples were drawn until a decline was observed in PHB production. To measure the cell dry weight, 5 ml ofculture sample was centrifuged at 8000 rpm for 10 minutes. The pellet was washed twice with distilled water and again centrifuged at 8000 rpm for 10 mins. The pellet was then dried at 80°C in hot air oven for about 12-24 hours until a constant weight was attained, cooled in a desiccator and weighed.

### Extraction by Sodium hypochlorite digestion

Extraction of PHB was done by Sodium hypochlorite digestion following the protocol mentioned. ^12^ The culture broth was centrifuged at 8000 rpm for 15 minutes. The pellet along with 10 ml Sodium hypochlorite solution was incubated at 50°C for 1 hour for lysis of the cells. The cell extract obtained was centrifuged at 12000 rpm for 30 minutes and then washed sequentially with distilled water, acetone and absolute ethanol. After washing, the pellet was dissolved in 10 ml chloroform and incubated at 50°C overnight and evaporated at room temperature. After evaporation, 10 ml of Sulphuric acid was added to it and placed in water bath for 10 minutes at 100°C. This converts the PHB into crotonic acid, which gives maximum absorption at 235 nm using sulphuric acid as blank.

### Spectrophotometric quantification by crotonic acid assay

PHB extracted from the above method was quantified by Crotonic acid assay.^13^ Crotonic acid powder was dissolved into 3 ml of Sulphuric acid and standard solution of 0.1μg of Crotonic acid/μl of Sulphuric acid was prepared. Working standards of 5, 10, 15, 20, 25, 30, 40, 50μg/3ml of Sulphuric acid were prepared. Blank was prepared by adding 3 ml of Sulphuric acid. The absorbance was measured at 235 nm. Standard graph of concentration v/s absorbance was prepared.

### Determination of PHB by GC-MS analysis

The polymer was prepared as described before and was characterized by GCMS analysis. ^12^ The extracted polymer was dissolved in 2mL of chloroform and then 2mL of methanol acidified with 3% (v/v) H_2_SO_4_ was added. The mixture was heated at 100°C for 3.5 hours for depolymerization and methanolysis of polyesters and 3µL was injected. The injection temperature was maintained at 260°C and column oven temperature was 100°C.

### Transmission Electron Microscopy analysis

For transmission electron microscopy (TEM) analysis, cells were centrifuged at 4,000 rpm for 20 minutes, following which the pellets were washed and then fixed in 4% (v/v) Glutaraldehyde and 1% Osmium tetroxide. The samples were dehydrated by passage through a series of increasing acetone concentrations (20, 40, 60, 80 and 100% acetone in water), embedded in epoxy resin, cut into ultra-thin 50nm sections and stained with uranyl acetate and lead citrate. The photomicrographs were captured at Cancer Institute, Chennai.

## ELECTROSPINNING PROCEDURE

The electrospinning apparatus was equipped with a high-voltage power supply with a maximum voltage of 50kV. The flow rate of the biopolymer solution was controlled by a precision pump to maintain a steady flow from the tip of the syringe needle. The electrospinning experiments were carried out under ambient conditions. The flow rate of the biopolymer solutions was fixed to 1ml/h and the applied voltage was 20 kV. The fibres were collected on a aluminium foil at a distance of 10 cm. ^9^

## ELECTROSPINNING OF POLMERIC BLENDS

### USING PVA AND PHB BLENDS

For production of PHB blends, About 15 % of PVA was dissolved in 90 ml of DMSO (Dimethyl sulphoxide) and 10 ml of PHB solution was added and the mixture was continuously stirred using a heating mantle, until a uniform blend was obtained.

### USING POLYVINYL ALCOHOL AND GELATIN-PHB FILM

For production of PHB blends, about 11 % of PVA was dissolved in 70 ml of DMSO (Dimethyl sulphoxide) and 30 ml of Gelatin-PHB solution was added and the mixture was continuously stirred using a heating mantle, until a uniform blend was obtained.

### USING PVA AND GELATIN-PHB FILM WIH ADDITION OF DMF

For production of PHB blends, About 11 % of PVA was dissolved in 70 ml of DMSO (Dimethyl sulphoxide) and 30 ml of Gelatin-PHB solution was added with addition of 200 µl of DMF (Dimethyl formaide) solution and the mixture was continuously stirred using a heating mantle, until a uniform blend was obtained.

## RESULTS

### Shake flask study and optimization of growth parameters

The bacterial isolate was grown in Minimal Salt Medium (MSM) to study the growth pattern of each strain and to determine the yield of PHB. It was shown that the maximum growth was attained on the third day with a maximum PHB yield of 49 mg/100 ml (0.49 g/ L). Figure 1 shows the growth curve of the selected bacterial isolate

**Figure 1:**
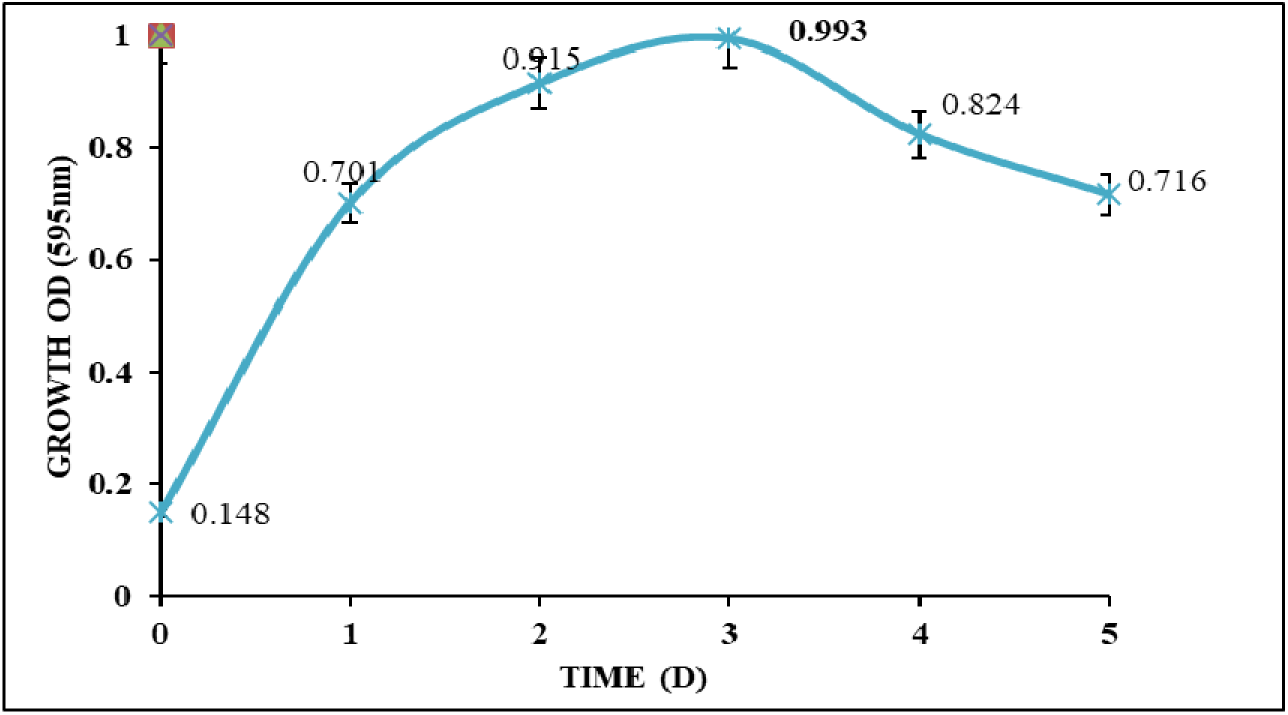
Growth curve of SNKVG 22.

**Figure 2.**
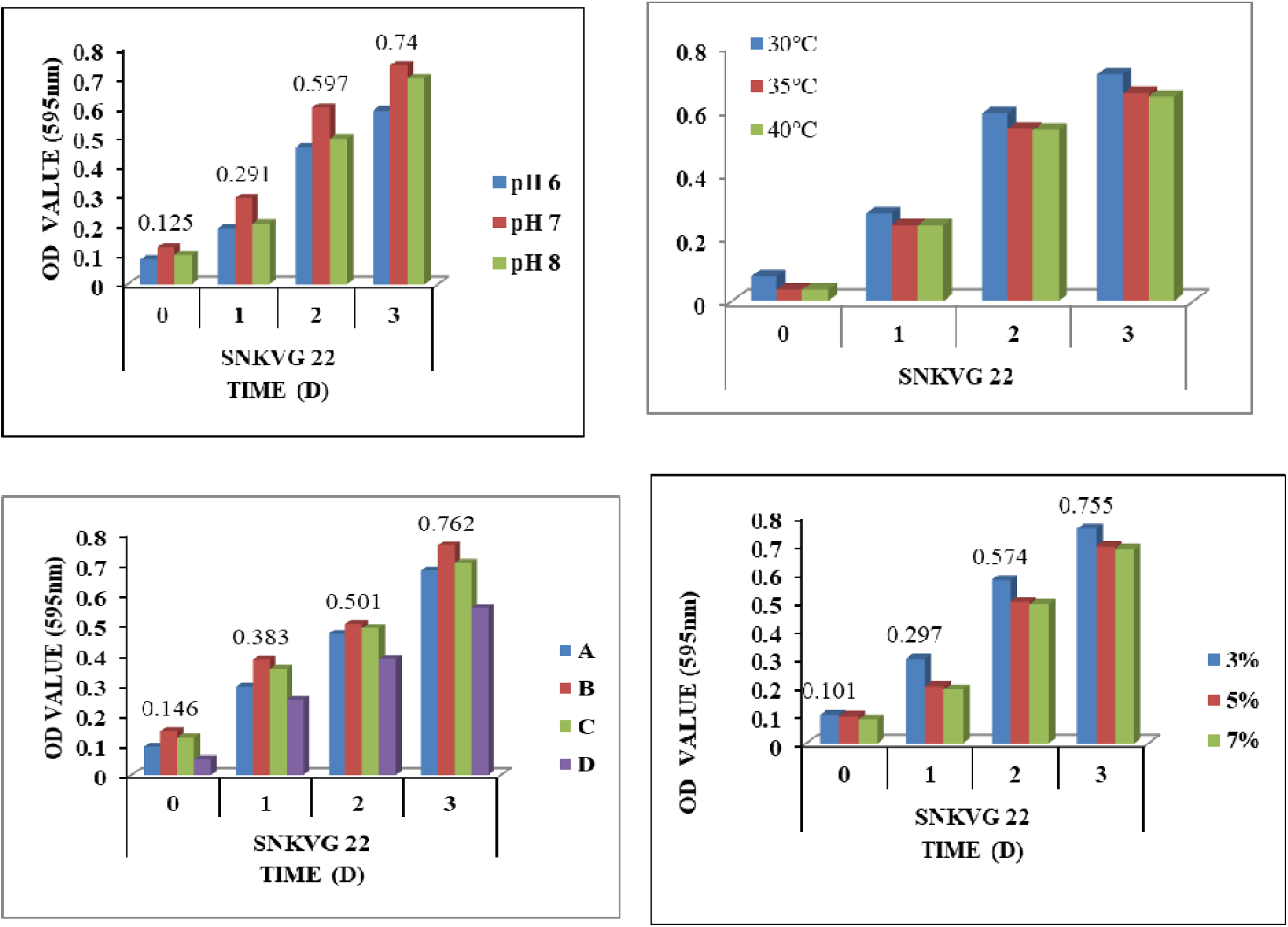
Optimization of growth conditions by the Bacterial strain.

### Reactor studies for large scale production of PHB

In the fermentation study of PHB production using molasses as an additional carbon substrate, it was observed that the maximum PHB production of 0.5 g/ L was attained at 24 hours of growth after which a decline in yield was observed. The temperature and pH was maintained throughout the fermentation process. The D.O. level was not controlled but was allowed to fall freely with 20 % saturation until equilibrium was established. Figure 3 shows the various parameters which were analysed during the fermenter study. It was seen that there was an increase in the Cell Dry Weight (CDW) as the specific growth rate increased at every 4 hours interval. At the end of 24 hours, the CDW was found to have increased with increase in the PHB production. The CDW still continued to increase slightly afterwards but the PHB production declined after 24 hours.

**Figure 3.**
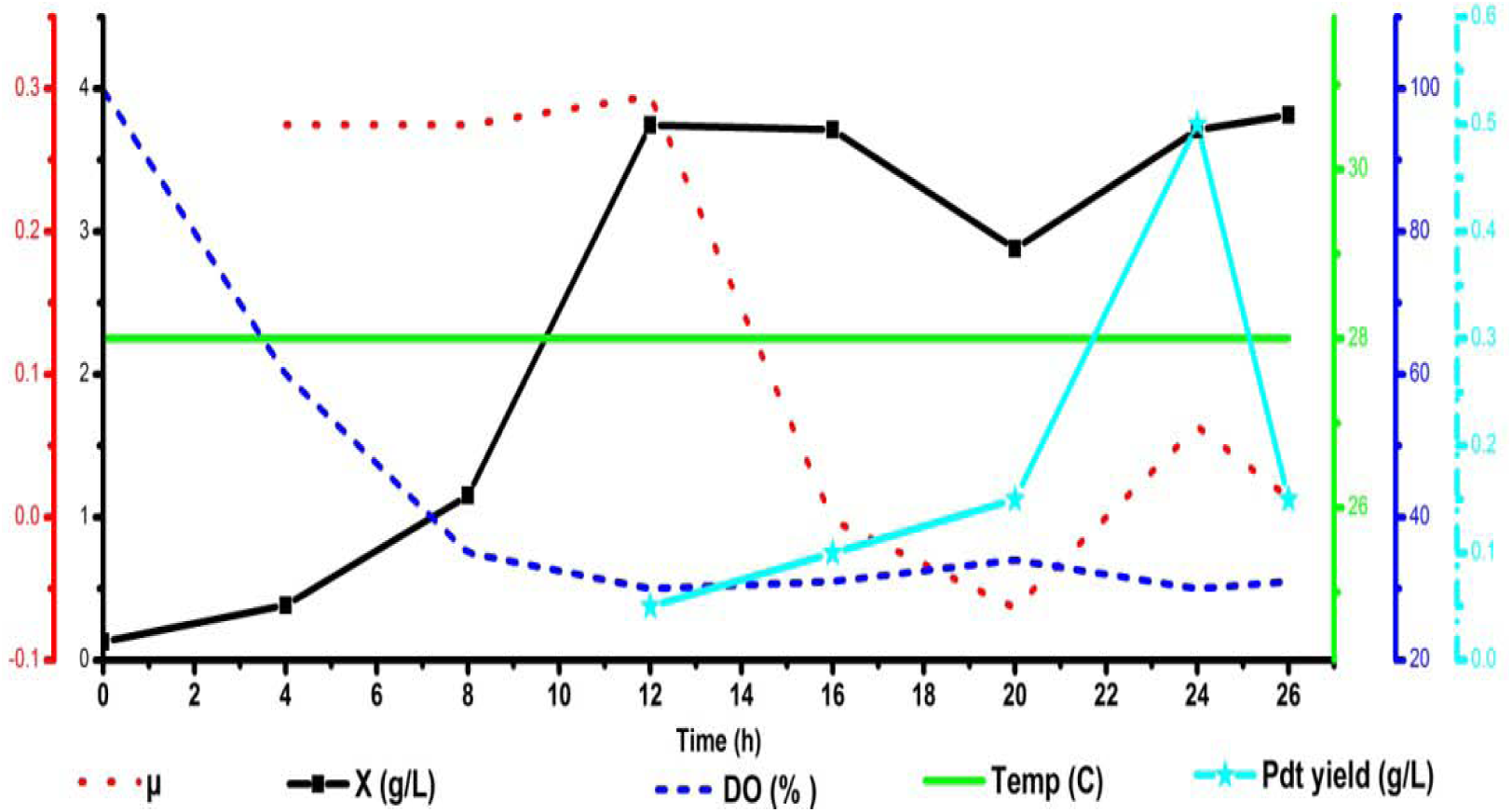
Various parameters observed during PHB production - specific growth rate (µ), Dissolved Oxygen (DO), Cell Dry Mass (X), Temperature and PHB yield.

## GC-MS ANALYSIS

In this study, the PHB was methanolysed in the presence of sulphuric acid and methanol, and it was then analyzed by GC-MS. The retention time was recorded and five different integration peaks at different retention times were observed. The retention times and ion fragment patterns of the peaks at 3.08 and 3.5 mins were identical to those of methylbenzylidene and propionic acid which are likely to be derivatives of PHB. The retention peaks at 5.48 and 5.95 mins correspond to dimethly ester groups of PHB. The integrated peak at 9.83 mins corresponds to methoxybenzylidine which promotes the liquid crystal formation in PHB. Figure 4 shows the integration peaks of the various compounds that were obtained at different retention times.

**Figure 4.**
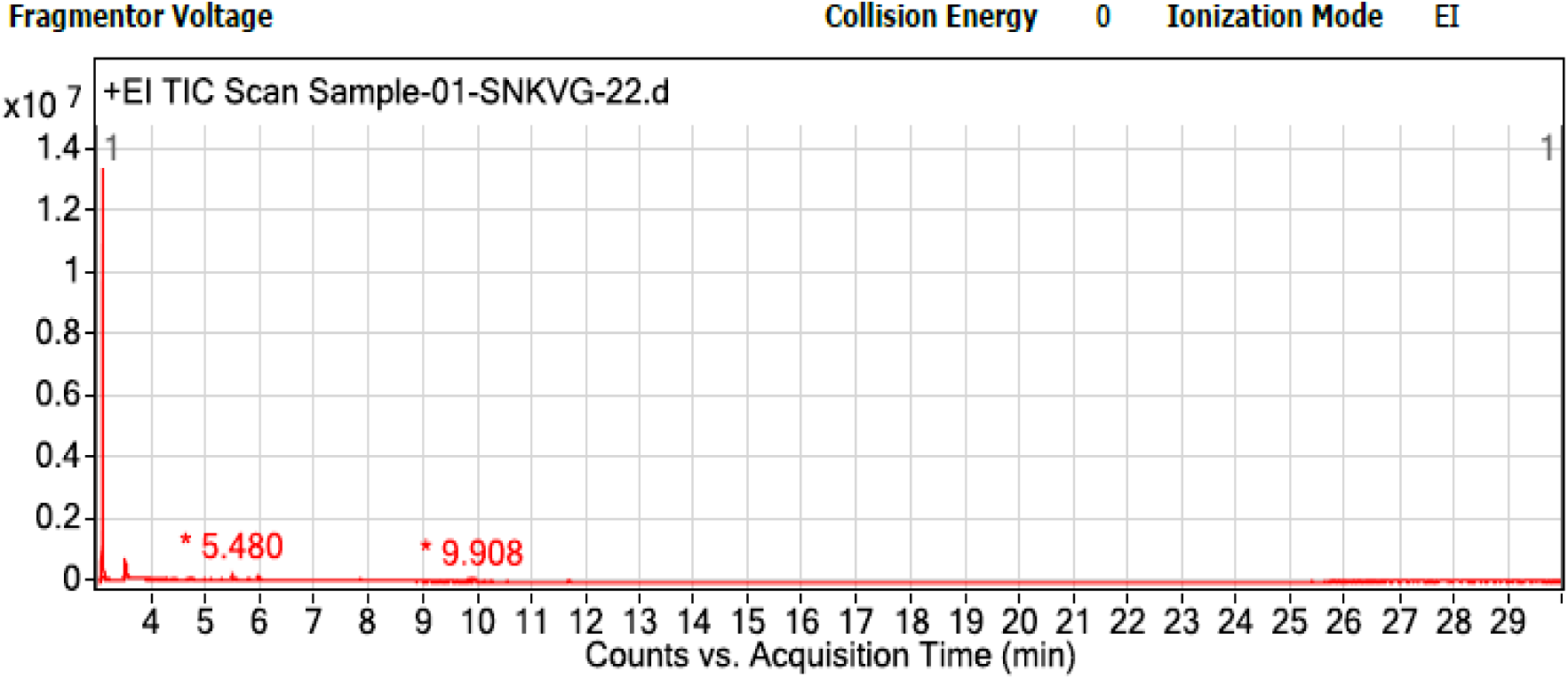
GCMS spectrum showing retention peaks in SNKVG-22.

## DETERMINATION OF PHB BY TEM ANALYSIS

Thin sections of the bacterial isolate SNKVG-22 were analyzed by TEM. The micrographs showed bacterial cells containing PHB granules in their cytoplasm. The PHB was seen as white inclusions against a black background. Figure 5 shows the PHB granule accumulated within the cytoplasm in the isolated bacterial strain.

**Figure 5.**
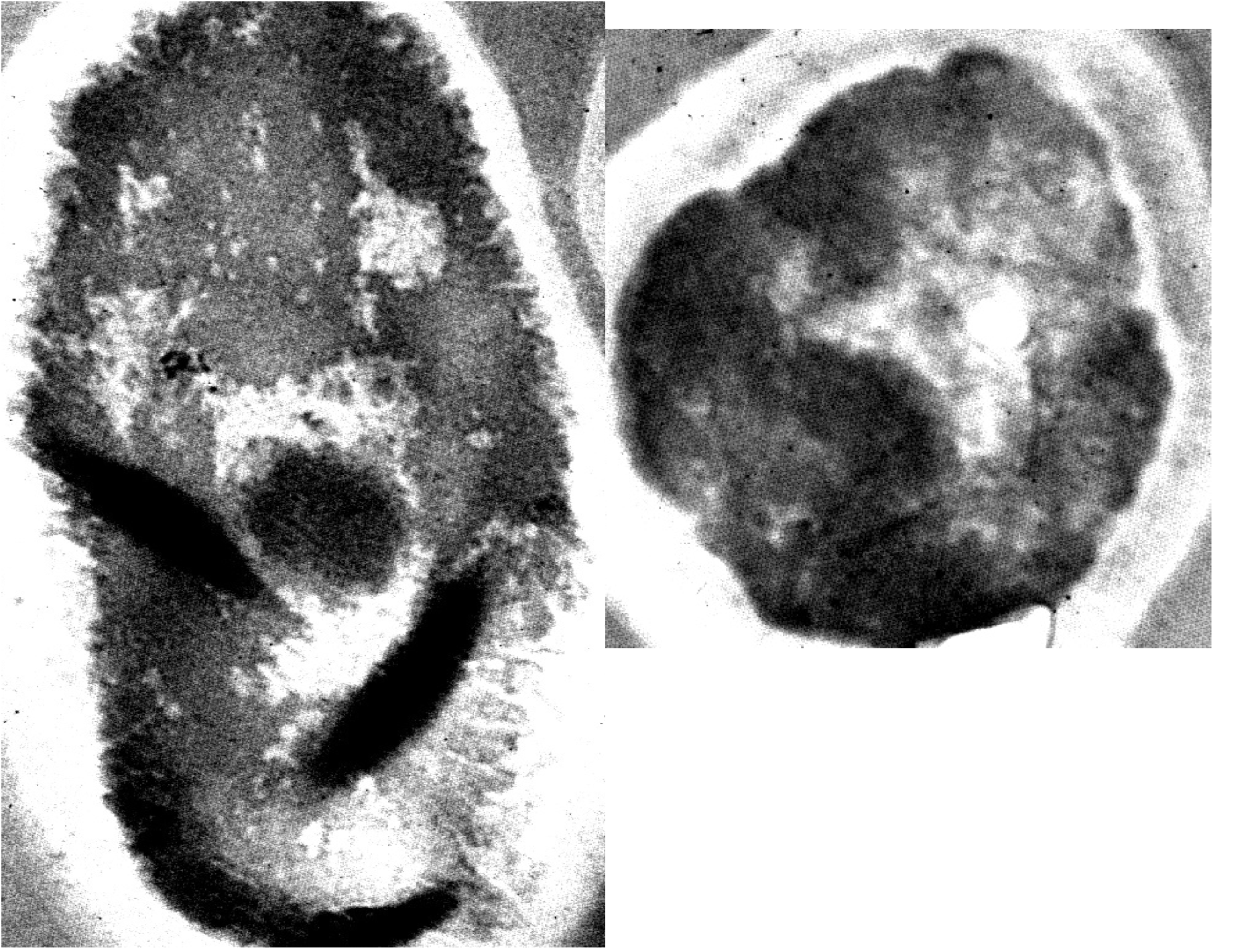
PHB granules seen in the cytoplasm of the bacterial strain SNKVG - 22 under TEM.

## APPLICATION OF PHB PRODUCED BY BACTERIAL STRAINS

### Gelatin-PHB films

A blend of gelatin and PHB were made into thin films. The Gelatin –PHB films were characterized by FT-IR Spectroscopy. The Gelatin concentration was initially optimized from 5 % to 20 % with 200 ml of distilled water and 100 µl of glycerol. Among optimization 20 % of Gelatin gave removable thin films. Later these films were used for electrospinning. Figure 6 shows the thin Gelatin-PHB films. Figure 7 and Figure 8 shows the FT-IR spectrum of the bacterial strains.

**Figure 6.**
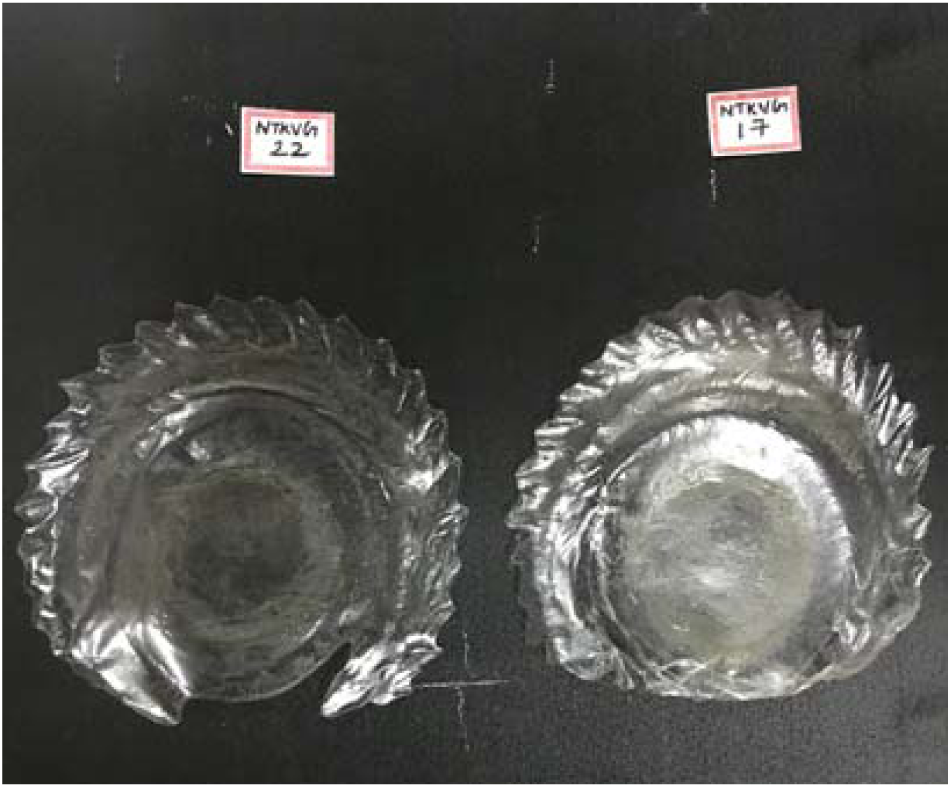
GELATIN-PHB FILMS.

**Figure 7.**
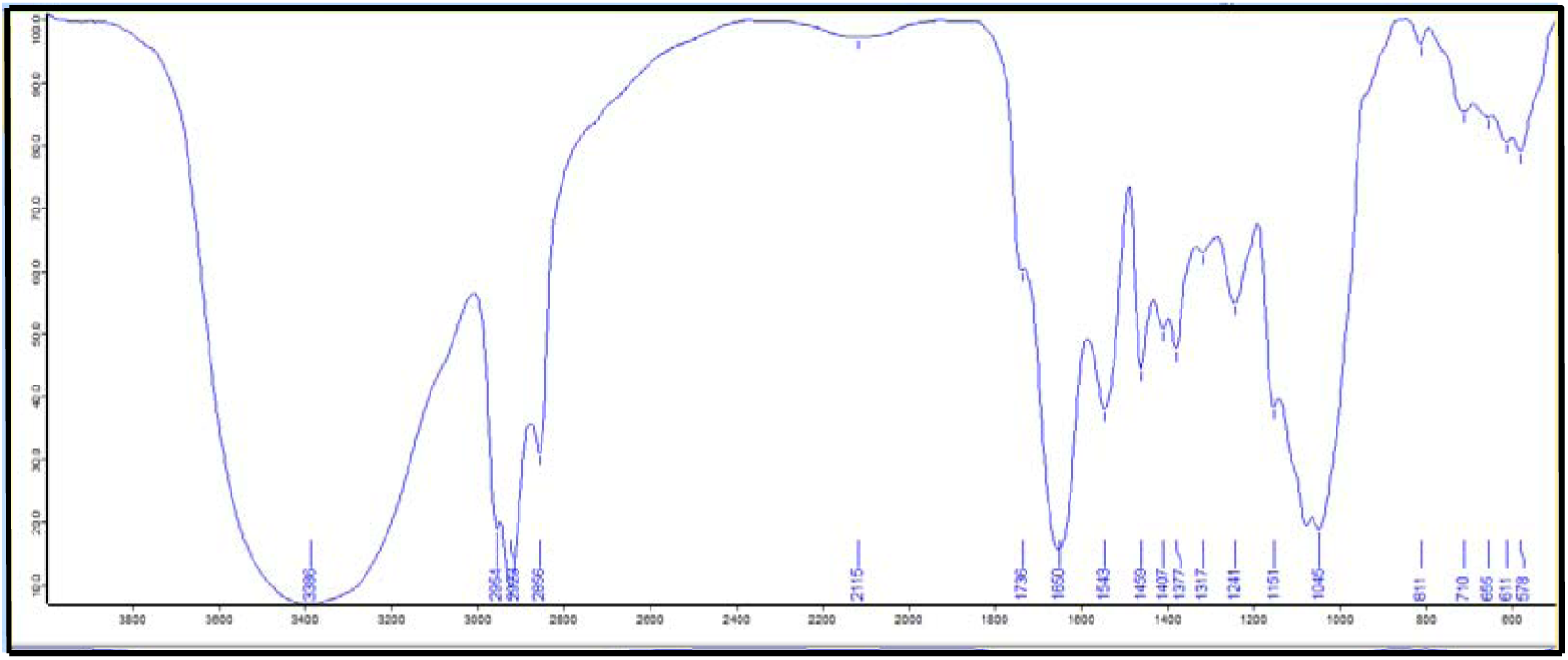
FTIR spectrum of bacterial isolate SNKVG 22 of GELATIN-PHB FILMS.

**Figure 8.**
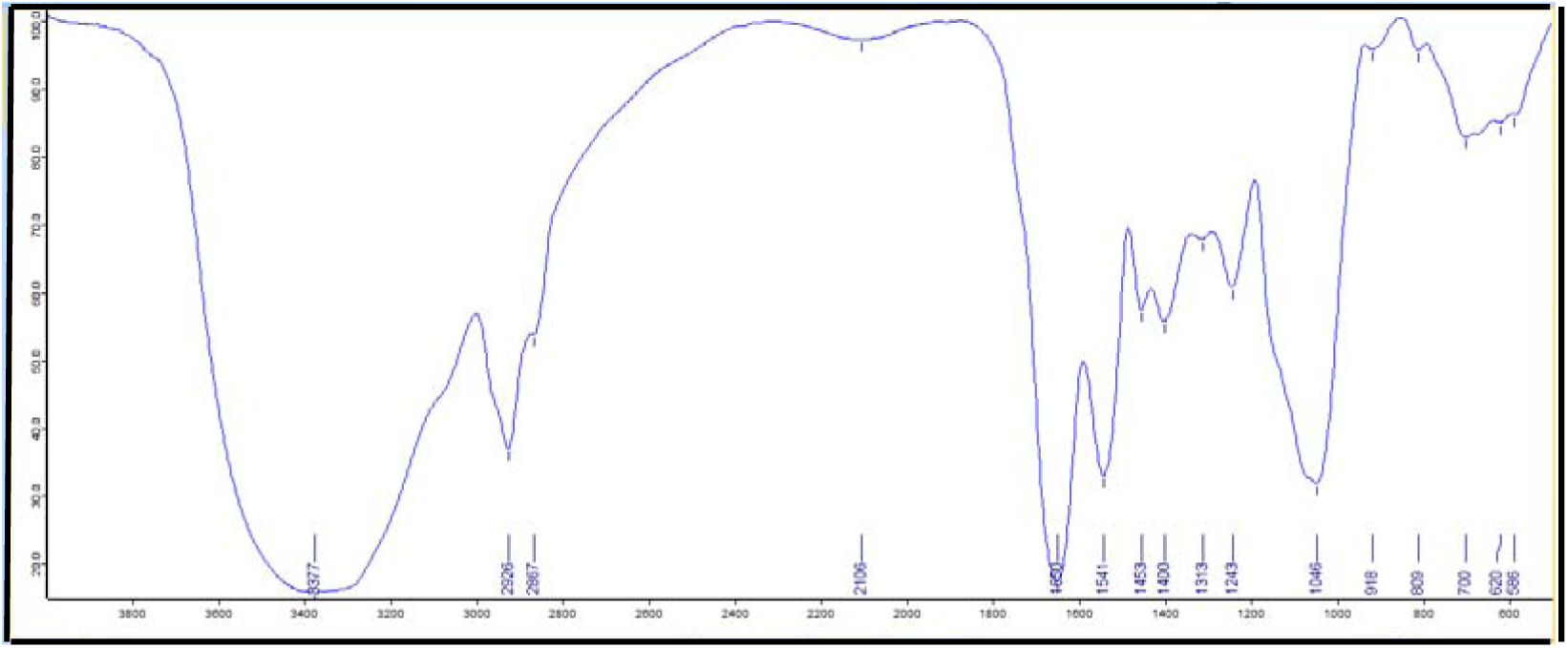
FTIR spectrum of bacterial isolate SNKVG 17 of GELATIN-PHB FILMS.

The two bacterial isolates were further tested for PHB production by FTIR Spectroscopy. The polymer was extracted as mentioned earlier from the two bacterial strains and was used for recording the IR Spectra in the range of 4000 – 500 cm^-1^. In the FTIR analysis results, the peaks at 3377 cm^-1^ indicates the stretching of H bond created by the terminal OH groups. The peak at the range of 1736 and 1650 cm^-1^ for the bacterial isolate which confirms the presence of alkyl ester groups with C = O stretching. The peak also showed absorption O – H stretching between 1453-1459 cm-^1^ for the isolates with variable and broad intensity. The peak in the range of 2923-2926 cm^-1^ are due to C – H stretching vibrations of methyl, methylene groups, whereas a series of peak between 1000 and 1300 cm^-1^ in all the isolates show stretching of C – O bond of the ester group. These prominent absorption bands confirm the structure of Poly-ß-hydroxybutyrate. Figure 7 and 8 shows the FT-IR spectrum of the bacterial isolates.

## ELECTROSPINNING OF PHB

In this study, the electrospinned sample was prepared by using 11 % PVA (polyvinyl alcohol), Gelatin and PHB as a blended and 15 % of PVA and PHB with DMSO (Dimethyl sulphoxide). The morphology of the electrospun fibers are loopy, indicating that there is a limited electrostatic repulsion force acting along the fiber during the spinning process. This is because in this case the electrospinning parameters were not optimized and will be done in future works. DMF (Dimethyl Formaide) (with higher dielectric constant and lower volatility) 100 µl was added to the solution, allowing to solve the above mentioned problems and resulting in stable and continuous processing of the fiber membranes. DMSO (Dimethyl sulphoxide) was used as a solvent to dissolve the PHB polymer and PVA, where the fibres obtained were non-uniform with water droplets and beads as DMSO has high boiling point the electrospun fibres obtained were disassociated. Figure 9 shows the Electrospinning apparatus used for the present study. Figure 10 shows the polymer membrane obtained by electrospinning using 15 % of PVA and PHB using DMSO (Dimethyl sulphoxide) as a solvent, showing the presence of gaps seen in the polymer membrane. Figure 11 shows Electrospinning fibres viewed under 40X magnification, which shows the presence of water droplets and uneven fibres. Figure 12 shows the SEM images of nanofibres obtained from electrospinning using 15 % of PVA and PHB using DMSO as a solvent at 15000 X magnification, the morphologies of the blend polymer were in sub-micron level diameter. Figure 13 shows the SEM images of nanofibres obtained from electrospinning using 15 % of PVA and PHB using DMSO as a solvent at 6000X magnification, the morphology of nanofibres were ranging from 82.9-141.9nm. Figure 14 shows the SEM images of nanofibres obtained from electrospinning using of 15 % of PVA and PHB using DMSO as a solvent at 30000X magnification, shows the nanobead ranging from 588.7 nm. Figure 15 shows the SEM images of the polymer membrane obtained from electrospining using 11 % PVA, Gelatin –PHB film. Figure 16 shows the electrospun nanofibres viewed under 40X magnification showing loopy fibres. Figure 17 SEM images of nanofibres obtained from electrospining using PVA, Gelatin-PHB films at 5000X magnification. Figure 18 shows SEM images of nanofibres obtained from electrospinning using PVA, Gelatin-PHB film at 60000X magnification, showing the morphology of the nanofibres ranging from 217-252 nm. Figure 19 SEM images of nanofibres obtained from electrospinning using PVA, Gelatin-PHB film at 15,000X magnification.

**Figure 9.**
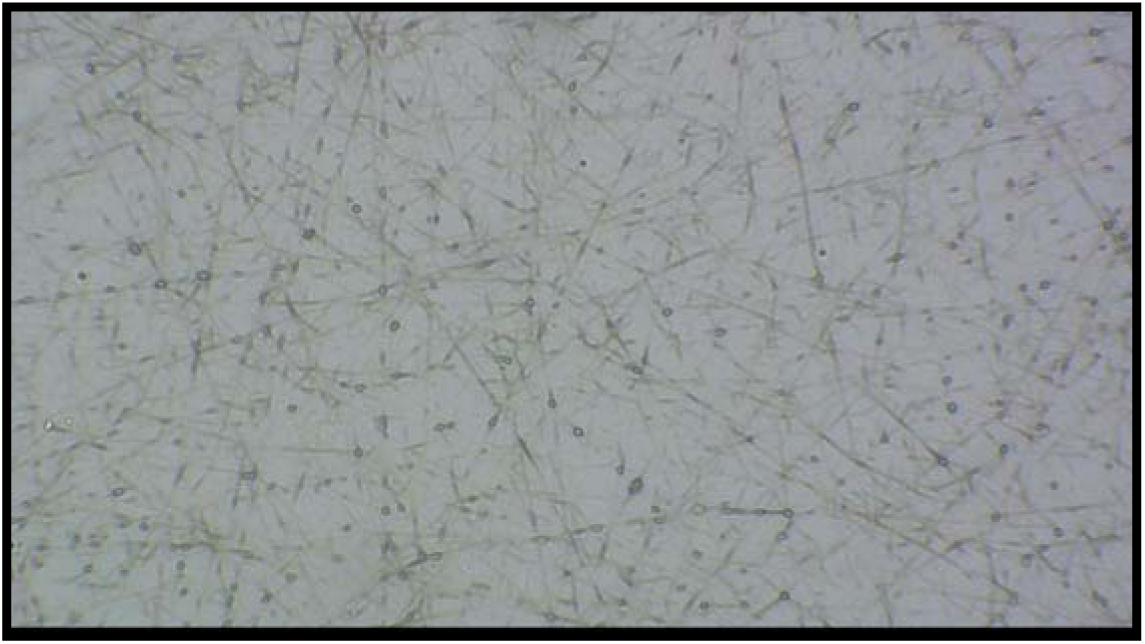
Electrospun fibers viewed under 40X Magnification.

**Figure 10.**
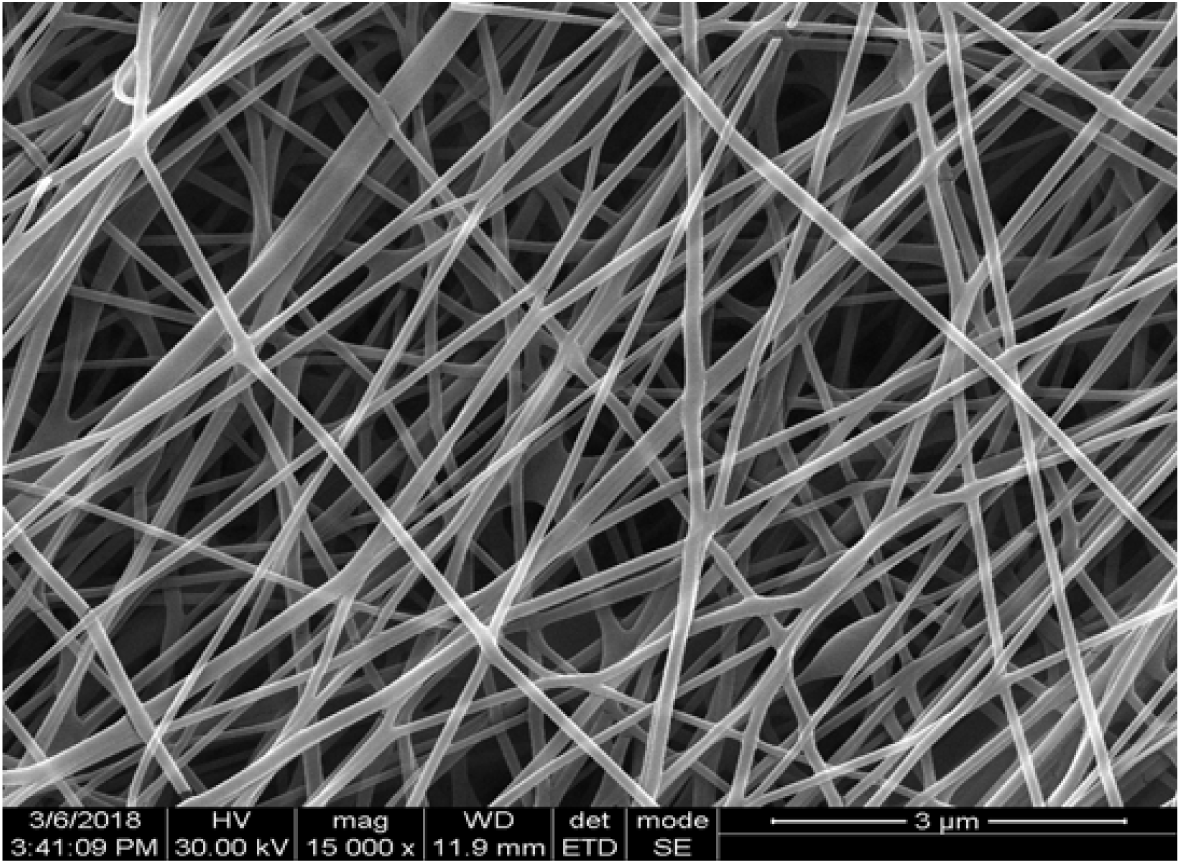
SEM images of nanofibres obtained from electrospinning using 15% of PVA and PHB using DMSO as a solvent at 15000X magnification.

**Figure 11.**
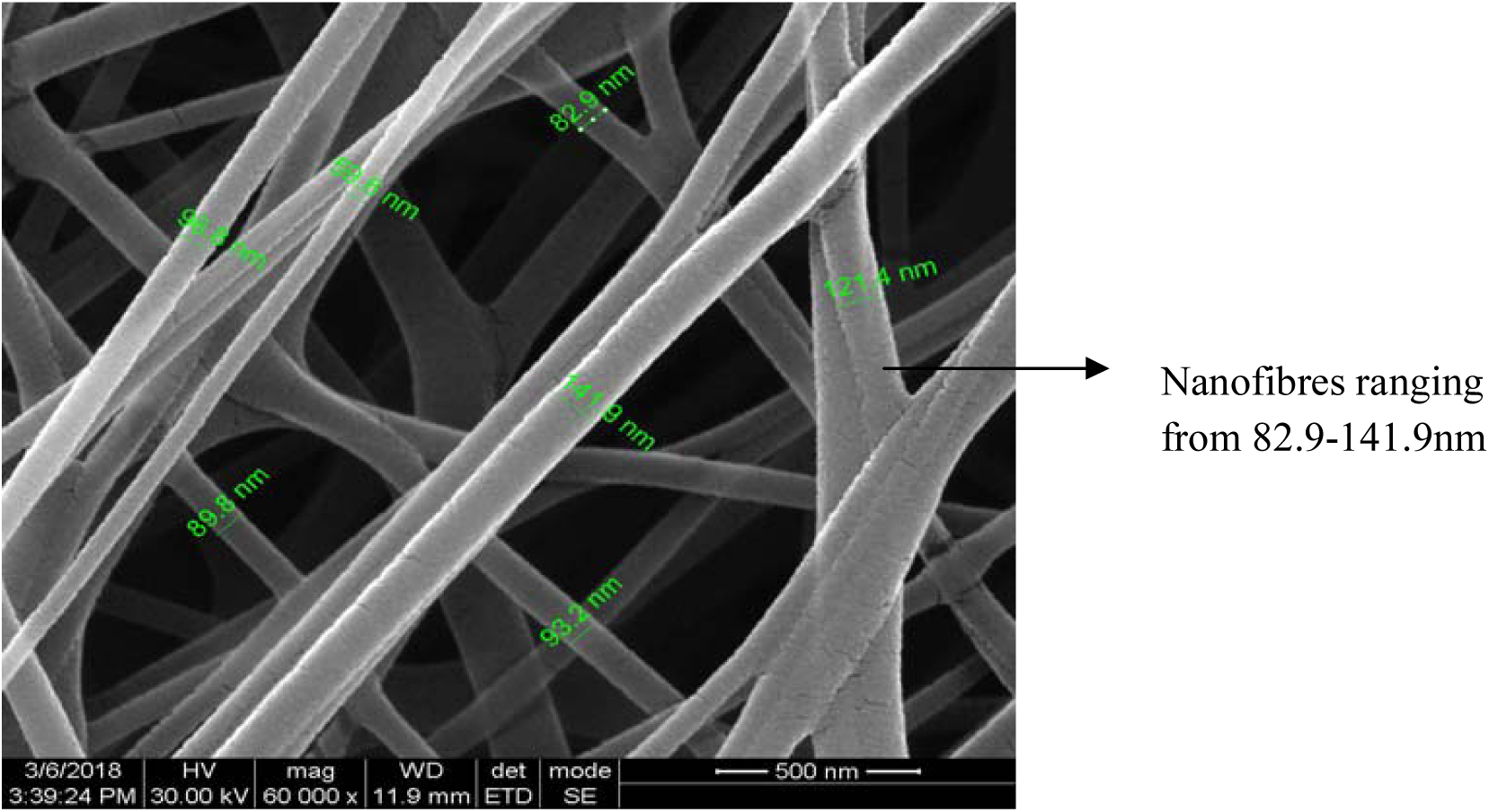
SEM images of nanofibres obtained from electrospinning using 15% of PVA and PHB using DMSO as a solvent at 60000X magnification.

**Figure 12.**
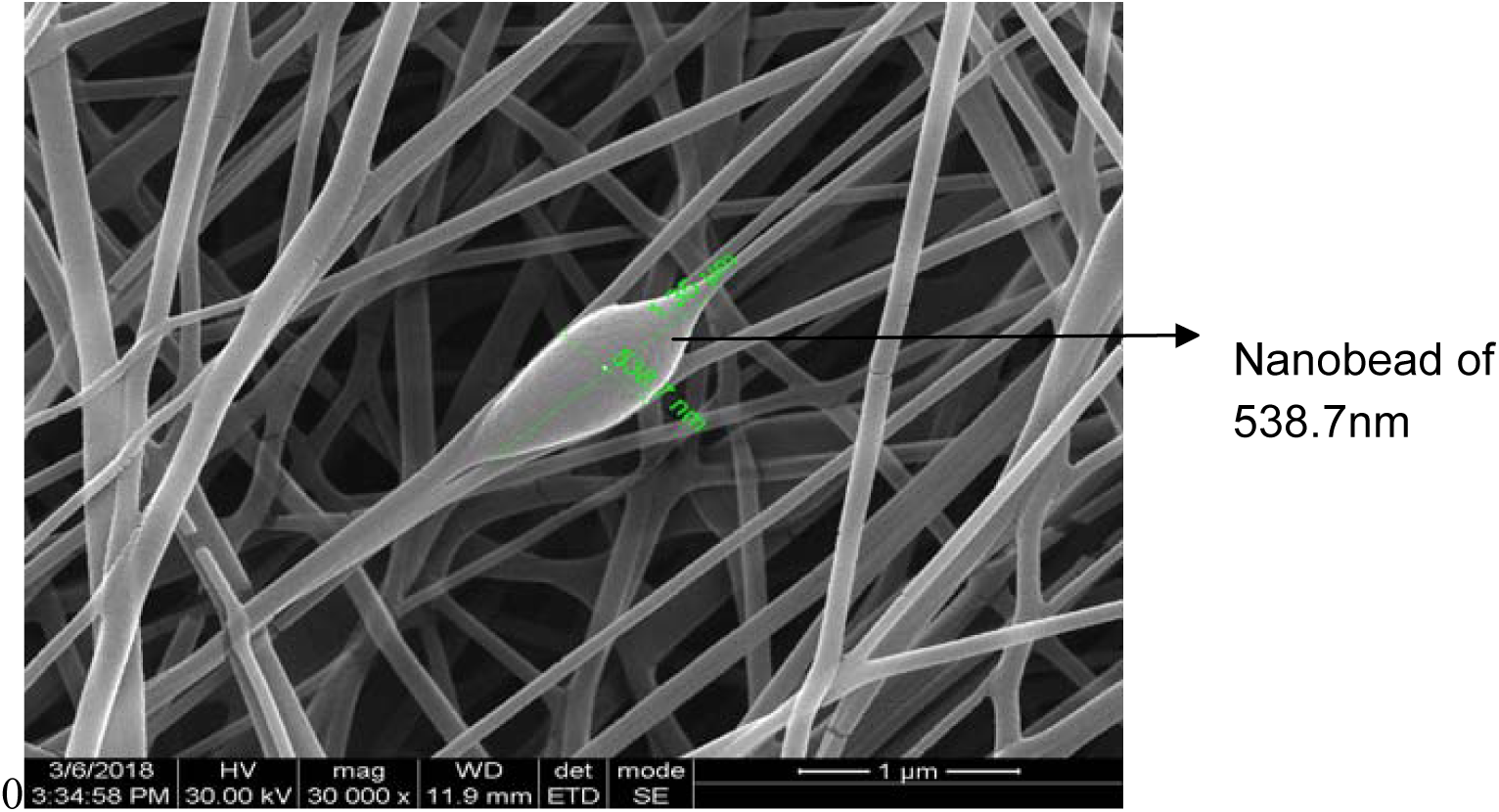
SEM images of nanofibres obtained from electrospinning using 15% of PVA and PHB using DMSO as a solvent at 30000X magnification.

## DISCUSSION

PHB and other PHAs are synthesized and deposited intracellularly in the form of granules and might amount up to 90% of the cellular dry weight (Schlegel*et al.* 1961). Various agro-industrial residues as carbon substrate for the production of PHB.^14^ Initially, in the presence of fructose, it was observed that the PHB yield and biomass concentration increased until 28^th^ hour of fermentation which was 25% and 1.1 g/ L respectively. On evaluation of various agro-industrial residues for PHB production, it was seen that jackfruit seed powder hydrolyzate gave best results with PHB production of 0.690 g/ L and PHB content in the cells was found to be 46%. Cane molasses showed some potential as a low-cost ingredient for industrial fermentation. Since cane molasses contains trace elements and vitamins such as thiamine, riboflavin, pyridoxine and niacinamide ^15,16^ it can also be used as a source of growth factors which maximized PHB production. In the present study, PHB was extracted from MSM using molasses as the carbon source along with glucose and the maximum PHB yield was found to be 0.5 g/ L at the end of 24^th^ hour with 3.7 g/ L of CDW.The decline in PHB production after 24 hours could be due to the fact that the microorganism could synthesize PHB until the sporulation stage and after that the remaining bacterial cells consume the PHB. The bacterial cells started utilizing PHB that was produced as their energy reservoir due to which the PHB yield declined but kept the cell dry weightalmost constant even after the 24^th^ hour.

GCMS helps in elucidating the structure of components. ^12^ identified the key components of PHB based on the retention time. The predominant peaks were noted at retention time of 11.6, 12.24, 13.6, 13.8 and 15.73 mins respectively which contained hexadecanoic acid, tetradecanoic acids and methyl esters which proved to be the monomer chains of the polyester family. In this study, retention times were recorded at 3.08, 3.5, 5.48, 5.95 and 9.9 mins which corresponded to the methyl ester groups and showed the presence of a number of monomers thereby proving the crude polymer production. Further purification of the polymers could help in characterizing the polymer in detail.

Transmission electron microscopy (TEM) analysis was done to reveal the accumulation of PHB within the bacterial cells. TEM analysis showed that the PHB granules can be seen as white inclusions which occupied most part of the cytoplasm of the bacterial cells against a dark background.^17^ In the present study, the cells showed small white inclusion bodies in the cytoplasm which were identified as PHB.

Bacterial plastic was prepared using PHB producing strains as biopolymer in 30 % concentration, 70 % of chemical polymer (sorbitol and gelatin) and glycerol as plasticizer. The commercial plastic and bacterial plastic was prepared using different ingredients with different concentration. ^18^ In this present study The Gelatin concentration was initially optimized from 5 % to 20 % with 200 ml of distilled water and 100 µl of glycerol 0.5 ml of gelatin solution and 5 ml of PHB solution was added to make thin GELATIN-PHB films. Among optimization tested 20 % of Gelatin gave removable thin films.

Electrospinning has garnered attention in recent years due to its potential for applications in various fields. Electrospinning fibres have shown great applicability for novel materials in tissue engineering, wound healing, bioactive molecules delivery as as sensor (Francisca *et al*., 2018). The parameters having influence on the morphology and properties of the electrospun fibers, Among the parameters related to the polymer solution, the most relevant are the nature of used solvent (dielectric properties, volatility, boiling point, and others), the solution concentration, that controls its viscosity, and the molecular weight of the polymer. For PHB, the solvent with high boiling point like DMSO (Dimethyl sulphoxide) allows a fast evaporation during the flight time. Full solvent evaporation occurs when the fiber reaches the grounded collector and therefore the feed rate does not have strong influence on fiber diameter (Daniela *et al*., 2013). In this present study the obtained electrospun fibers were spinned with 15 % and 11 % PVA with PHB and Gelatin. The morphology of the nanofibres were initially disassociated and there was presence of water droplets, after optimisation with decreasing the PVA and solvent concentration the nanofibres obtained were loopy indicating that there is electrostatic repulsion along the spinning process, also the influence of solvent like DMSO with the polymer let to the evaporation and formation of bead in the fibres. Later optimization was done with other polymer blends with addition of DMF (Dimethyl Formaide), in order to obtain thin loopy fibres.

